# Urban population structure and dispersal of an Australian mosquito (*Aedes notoscriptus*) involved in disease transmission

**DOI:** 10.1101/2022.01.18.476837

**Authors:** Véronique Paris, Rahul Rane, Peter Mee, Stacey Lynch, Ary A Hoffmann, Thomas L Schmidt

**Affiliations:** Pest and Environmental Adaptation Research Group, School of BioSciences, Bio21 Institute, University of Melbourne, Parkville, VIC, Australia; CSIRO Black Mountain Laboratories, Clunies Ross Street, Canberra, ACT, Australia; Agriculture Victoria Research, AgriBio Centre for AgriBiosciences, Bundoora, VIC, Australia

**Keywords:** Dispersal, mosquito, intergenerational dispersal, Aedes notoscriptus, kinship

## Abstract

Dispersal is a critical factor in designing successful pest control measures as it determines the rate of movement across target control areas and influences the risk of human exposure to the species and its pathogens. Here we used a fine scale spatial population genomic approach to investigate the dispersal ecology and population structure of *Aedes notoscriptus*, an important disease transmitting mosquito, on the Mornington Peninsula near Melbourne, Australia. The species is suspected to be involved in the transmission of *Mycobacterium ulcerans*, the bacterium that causes Buruli ulcer, in this area. We sampled and reared *Ae. notoscriptus* eggs at two time points from 170 traps up to 5 km apart and generated genomic data from 240 individuals. We also produced a draft genome assembly from a laboratory colony established from mosquitoes sampled near the study area. We found low genetic structure (Fst) and high coancestry throughout the study region. Using genetic data to identify close kin dyads, we found that mosquitoes had moved distances of >1km within a generation, which is further than previously described for this species. A spatial autocorrelation analysis of genetic distances indicated genetic dissimilarity at >4 km separation, a fourfold higher distance than for a comparable population of the dengue mosquito, *Ae. aegypti*, from Cairns, Australia. These findings point to high mobility of *Ae. notoscriptus*, highlighting the challenges of localized intervention strategies targeting this species. Further sampling within the same area at two time points 6 and 12 months after initial sampling showed that egg counts were relatively consistent across time, and that spatial variation in egg counts covaried with spatial variation in Wright’s neighbourhood size (NS). As NS increases linearly with population density, egg counts may be useful for estimating relative density in *Ae. notoscriptus*. The results highlight the importance of acquiring species-specific data when planning control measures.

## 1. Introduction

For insects that transmit disease, dispersal is a critical ecological characteristic when planning management strategies. Dispersal will influence the movement into and out of controlled areas as well as the extent of how the human, animal, or plant populations will be exposed to disease. Intrinsic and extrinsic factors such as dispersal barriers (natural or anthropogenetic) (Goldberg & Lande 2015; Schmidt *et al*. 2022), urbanisation (Johnson & Munshi-South 2017), and habitat fragmentation (Doak *et al*. 1992) can all influence insect dispersal. Once dispersal patterns are understood they can be used to enhance the effectiveness of insect pest control strategies. For example, the larval dispersal pattern of the fruit orchard pest *Operophtera brumata* is used to indicate when farmers should implement control measures to mitigate the damage to plants while decreasing the use of insecticides (Edland 1983; Jeger 1999). Dispersal studies focusing on the primary vector of dengue, *Aedes aegypti*, have shown how the dispersal of the vector influences the spread and distribution of the disease, which shaped the course of dengue control measures around the globe (Liew & Curtis, 2004; Harrington *et al*. 2005; Carvajal *et al*. 2020).

Mosquitoes of the genus *Aedes* are important vectors of human and animal diseases. Their control traditionally involves using insecticides (McCarroll *et al*. 2000), the removal of larval habitats, and the deployment of traps to reduce population sizes (Juarez *et al*. 2021). Novel interventions are now also applied, including the release of mosquitoes carrying novel *Wolbachia* bacteria (Hoffmann *et al*. 2011) or genetically modified mosquitoes (Harris *et al*.2012) to reduce or manipulate populations. All these strategies rely on knowledge about the dispersal of the target species, which will influence the effectiveness and long-term success of control programs. For instance, in *Ae. aegypti Wolbachia* stability and invasion into surrounding areas depends on mosquito movement across a local area (Schmidt *et al*. 2018).

Numerous methods have been deployed to estimate dispersal in *Aedes* mosquitoes. Mark-Release-Recapture (MRR) studies are commonly used (Reiter *et al*. 1995; Honório *et al*.2003) but can come with drawbacks if applied to small organisms like mosquitoes. They can be labour intensive, recapture rates can be low, and mosquitoes can be impacted by the marking method. Recent approaches try to overcome those difficulties by expanding MRR approaches to incorporate genetic inferences of close kin, functioning on the basis that an individual’s genotype can be considered a “recapture” of the genotypes of its relatives. This close kin mark-recapture (CKMR) framework has mainly been used to investigate abundance in large populations and to assess migration between populations (Palsøll *et al*. 2010). Recently, the use of genome-wide sequence data expands CKMR, which detects dispersal by assigning dyads to kinship categories across multiple orders of kinship and then using the spatial distribution of these kin to reveal past dispersal over fine temporal scales (Schmidt *et al*. 2018; 2021; 2022, Combs *et al*. 2018; Fountain *et al*. 2018; Jasper *et al*. 2019; Trense *et al*. 2021; Aguillon *et al*. 2017). These recent studies revealed the power of novel genetic approaches to estimate individual dispersal in small organisms like *Aedes* mosquitoes. They also demonstrate an opportunity to use the genetic data acquired for additional analyses that go beyond dispersal, such as investigations of population structure and dynamics (e.g., neighbourhood size (Jasper *et al*. 2019); level of gene flow (Combs *et al*. 2018)).

Here we use spatial population genomics to investigate the population structure and fine-scale dispersal of *Ae. notoscriptus* (the Australian backyard mosquito), a container breeding mosquito native to mainland Australia and Tasmania and invasive in the Torres Strait Islands, New Zealand, New Guinea, New Caledonia, Indonesia (Dobrotworsky 1965; Lee *et al*. 1987; Sunahara & Mogi 2004) and more recently in California, USA (Metzger *et al*. 2021). This species transmits arboviruses (e.g., Ross River virus & Barmah Forest virus) and dog heartworm (Russell & Geary 1992; Doggett & Russell 1997; Watson & Kay 1999) and is also suspected to be a significant vector of *Mycobacterium ulcerans*, the bacterium that cases Buruli ulcer (BU) (Wallace *et al*. 2017; Mee *et al*. unpublished data). Recent molecular studies have indicated several genetically different but morphologically similar populations of *Ae. notoscriptus* distributed across Australia (Endersby *et al*. 2017). In Victoria, two main clades occur in sympatry, and it is yet to be determined if different clades of *Ae. notoscriptus* differ in ecological characteristics such as habitat, feeding, and mating preferences, which can influence dispersal and the species’ ability to transmit diseases. Though this mosquito is an important vector, threatening human and animal health, little research focuses on its population dynamics in the field, which makes risk calculations and planning of interventions difficult. We present a draft reference genome for *Ae. notoscriptus* and use a genomic approach to investigate individual-level dispersal and spatial genetic structure of *Ae. notoscriptus* at the Mornington Peninsula, where BU cases have clustered in recent years (https://www.mornpen.vic.gov.au/Community-Services/Health-Wellbeing/Health-Safety/Buruli-Ulcer). We also calculate Wright’s neighbourhood size (NS: Wright 1946) estimates throughout the study area, and compare these to egg counts to determine whether egg count data sufficiently reflect local *Ae. notoscriptus* abundance.

## 2. Materials & Methods

### 2.1 Reference genome assembly of *Aedes notoscriptus*

#### 2.1.1 DNA extraction and sequencing

We extracted DNA of *Ae. notoscriptus* from a laboratory culture sampled initially in Cheltenham, Victoria, Australia, in 2018. DNA extraction was carried out using Qiagen genomic tip 20/g following manufacturer’s recommendations. An Illumina PCR-free library was then constructed using IDT Lotus kits and a fragment size of 250bp. The library was then sequenced on Illumina HiSeqX 150bp PE to obtain 136.22 Gb raw sequence data.

#### 2.1.2 Genome assembly

The raw paired-end data was filtered to remove adapter sequences using *TrimGalore* v0.6.6 resulting in 135 Gb of clean data. This was then assembled using *MaSuRCA* v3.4.0 (Zimin *et al*. 2013) and *SparseAssembler* v1 (Ye *et al*. 2012) to obtain high fidelity contigs, which were then merged using *flye* v2.7 (Kolmogorov *et al*. 2019). This merged assembly was then deduplicated using *purge_dups* v1.2.4 (Guan *et al*. 2020) to create the draft assembly for analysis spanning 1.55 Gb with an N50 of 3893 bp.

### 2.2 Fine scale population structure and dispersal

#### 2.2.1 Sampling of *Ae. notoscriptus*

In February 2019, we sampled *Ae. notoscriptus* from four sites at the Mornington Peninsula: Sorrento (‘north-west’), Blairgowrie (‘central’) and Rye (‘north-east’ and ‘south-east’) (Figure 1).

**Figure 1.**
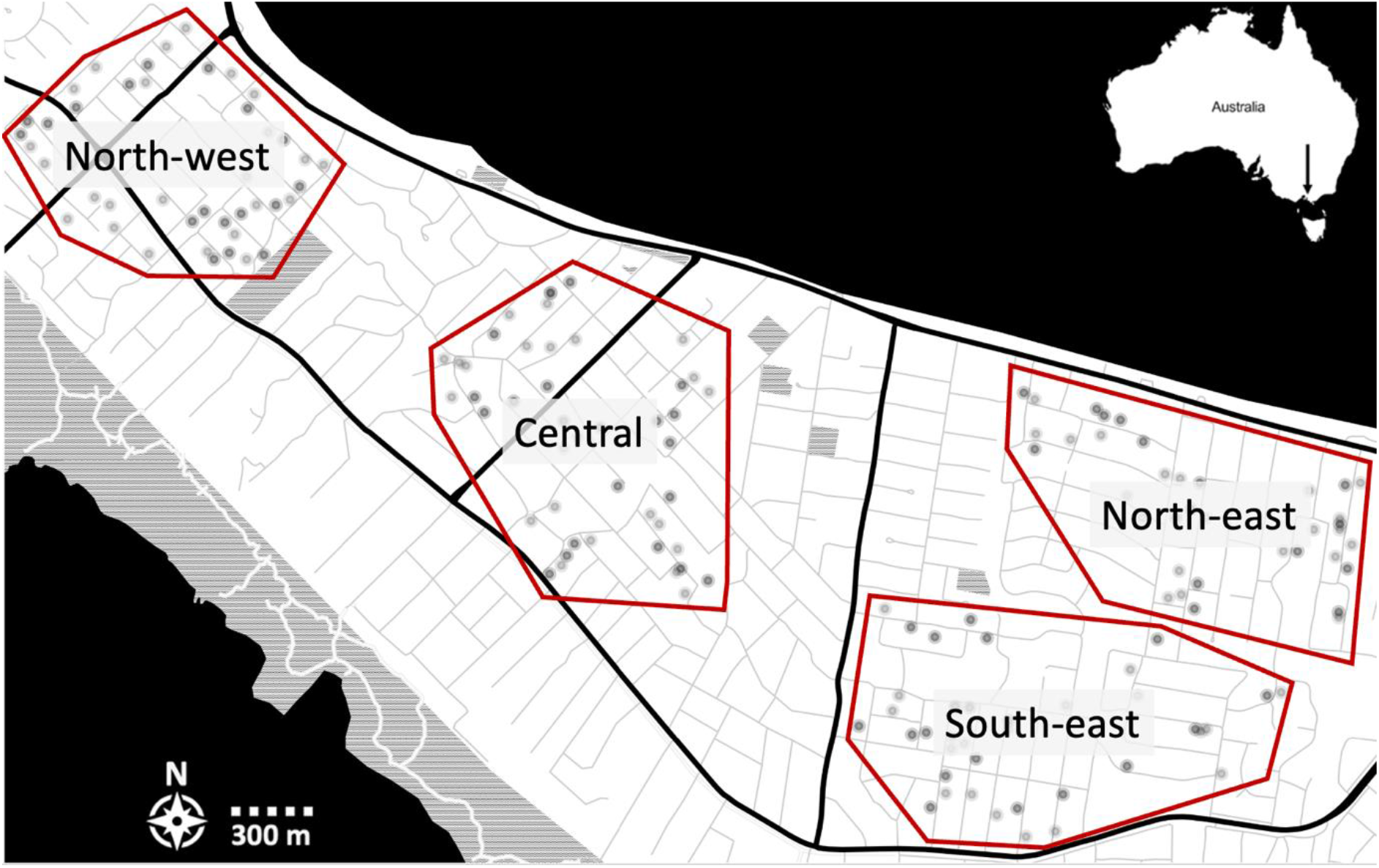
Sampling sites at the Mornington Peninsula. ‘north-west’, ‘central’, ‘south-east’ and ‘north-east’ sites from left to right indicated by red rectangles. Transparent black circles indicate trap locations in each sampling site. Trap placement is visualized in more detail in Figure 4.

Sampling used 340 oviposition traps (~90 traps per sampling site) consisting of a black plastic bucket half-filled with water and containing several alfalfa pellets to attract gravid *Ae. notoscriptus* (Ritchie 2001). The average distance between traps was 1046 m (ranging from 16 m to 5480 m). A ~15 cm long strip of red felt extending into the water provided an oviposition substrate. We collected felts after 7 and again after 14 days, which were partially dried to encourage hatching. Three days after collection, we hatched eggs in 500 mL reverse osmosis (RO) water containing 2-3 TetraMin tropical fish food tablets (Tetra, Melle, Germany). If no larvae hatched, felts were re-dried for three days and the hatching process repeated. We replaced water and food as appropriate. Emerging virgin adults were transferred into absolute ethanol and stored at −20°C until DNA extraction. We selected 240 individual mosquitoes for DNA sequencing, covering 170 different trap locations. To maximise spatial distribution, we prioritised samples collected from different traps over samples from different weeks. Care was taken to only include one individual per felt to avoid the detection of close kin sampled from the same felt.

#### 2.2.2 DNA extraction and library preparation

Using keys from Dobrotworsky (1965) we morphologically identified *Ae. notoscriptus* mosquitoes and extracted DNA from individuals using either Qiagen DNeasy Blood & Tissue Kits (Qiagen, Hilden, Germany) or Roche High Pure PCR Template Preparation Kits (Roche Molecular Systems, Inc., Pleasanton, CA, USA) following the manufacturers’ instructions. We prepared double-digest restriction site-associated DNA sequencing (ddRADseq) libraries following the method for *Ae. aegypti* (Rašić *et al*. 2014). We started with an initial digestion of 10 ng of genomic DNA, ten units each of MluCI and NlaIII restriction enzymes, NEB CutSmart buffer (New England Biolabs, Beverly, MA, USA), and water. Digestions were run for 3 hours at 37 °C with no heat kill step, and the products were cleaned with paramagnetic beads. Modified Illumina P1 and P2 adapters were ligated onto cleaned digestions overnight at 16 °C with 1,000 units of T4 ligase (New England Biolabs, Beverly, MA, USA), followed by a 10-minute heat-deactivation step at 65 C. We performed size selection using a Pippin-Prep 2% gel cassette (Sage Sciences, Beverly, MA) to retain DNA fragments of 350 – 450 bp.

The size selected libraries were amplified by PCR, using 1 μL of size-selected DNA, 5 μL of Phusion High Fidelity 2× Master mix (New England Biolabs, Beverly MA, USA) and 2 μL of 10 μM standard Illumina P1 and P2 primers. These were run for 12 PCR cycles, then cleaned and concentrated using 0.8x paramagnetic beads. Each ddRAD library contained 24 mosquitoes, and each was sequenced on a single sequencing lane using 150 bp chemistry. Libraries were sequenced paired-end at GeneWiz, Inc (Suzhou, China) using a HiSeq 4000 (Illumina, California, USA).

#### 2.2.3 Data processing

We used the *Process radtags* program in *Stacks* v2.0 (Catchen *et al*. 2013) to demultiplex sequence reads. Low-quality reads were discarded using a 15 bp sliding window if the average Phred score dropped below 20. We used *Bowtie* v2.0 (Langmead & Salzberg 2012) to align reads to the *Ae. notoscriptus* reference genome assembly (described in 2.1), using —*very-sensitive* alignment settings. We filtered all alignments to paired reads that aligned concordantly, requiring the two paired reads to align to the same contig to avoid multi-mapping using *Samtools* (Danecek *et al*. 2021). *Stacks Ref_map* program was used to build *Stacks* catalogs, from which genotypes were called at a 0.05 significance level and *--min-mapq* 15 to filter any remaining multi-mapped reads. We generated VCF files for the catalog with the *Stacks* program *Populations* (Catchen *et al*., 2013). Single nucleotide polymorphisms (SNPs) were required to be scored in ≥ 90% of mosquitoes, with a minor allele count of ≥ 3 (*-r* 0.90 *–mac* 3 *--vcf*). *Beagle* v4.1 (Browning & Browning 2016) was used to impute and phase the dataset in a 50,000 bp sliding window with 3,000 bp overlap. Finally, *vcftools* thinned SNPs so that no two SNPs are at 500 bp distance to each other (*--thin* 500). After filtering, we retained 11,091 SNPs.

#### 2.2.4 Genetic diversity and local population structure

*Populations* was used to calculate pairwise FST values between the four sampling sites. We tested for isolation by distance (IBD) between all samples and within each of the four sampling sites using the *mantel.randtest* function in the *R* package *ade4* (Dray & Dufour 2007). We also tested for IBD in the dataset after removing pairs of individuals identified as close kin, as close kin can drive IBD patterns at sufficiently fine scales (Aguillon *et al*. 2017). The simple Mantel tests analysed pairwise genetic distance matrices and the natural logarithm of Haversine pairwise geographic distance, employing 9,999 permutations and Bonferroni correction to assess statistical significance. Rousset’s *a* (Rousset 2000) provided genetic distances, calculated in *SPAGeDi* (Hardy & Vekemans 2002). Additionally, we used the pairwise genetic distance and geographical distance matrices to measure spatial autocorrelation using the *mgram* function in the *R* package *ecodist* (Goslee & Urban 2007) to build correlograms.

To contextualise these spatial genetic structure results, we compared results for *Ae. notoscriptus* with an *Ae. aegypti* population from Cairns, Australia, sampled in 2015 using similar protocols and with a similar spatial distribution of traps (Schmidt *et al*. 2018). This *Ae. aegypti* dataset contained both individuals carrying a *Wolbachia* infection (*w*Mel) from recent releases in the area and wildtype individuals without the infection (WT). We analysed these separately to avoid bias. We down sampled each *Ae. aegypti* dataset to only include one sample per trap to achieve maximum comparability to the *Ae. notoscriptus* dataset.

To further investigate the local population structure of *Ae. notoscriptus* and coancestry between individuals of the different sampling sites, we used the program *fineRADstructure* (Malinsky *et al*. 2018), which we ran with default settings.

#### 2.2.5 Local dispersal estimates of individuals

We investigated the association between pairwise kinship and distance to infer specific dispersal of the parental generation, treating separation distances between pairs of kin as representative of past dispersal events. The distances between full-siblings result from the mother’s oviposition dispersal and therefore represent the direct dispersal of a single individual female between two traps. Female *Aedes* mosquitoes usually only mate once, while males mate with several females (Christophers, 1960). Hence, half-sibling separation distances result from past mating dispersal of their father and the dispersal of their mothers for host-seeking and oviposition. First cousins are separated through the ovipositional dispersal of their grandmother and the premating dispersal of each parent, plus the post-mating dispersal and ovipositional dispersal of each mother (Jasper *et al*. 2019).

We generated kinship coefficients using *PC-Relate* (Conomos *et al*. 2016), which conditions the data with principal components (PCs) to control for genetic structure. We generated kinship coefficients for all dyads following different conditioning treatments, ranging from 2PCs to 30 PCs. The PC plots revealed that 4PCs conditioned the data the best, with no structure observed in additional PCs (Figure S1). We defined kinship classes with full-siblings kin ≥ 0.1875 and half-siblings ≥ 0.09375. To determine the lower bound to define the category of first cousins, we reviewed a scatterplot of the estimated kinship coefficients and estimated probabilities of sharing zero alleles IBD (k0) (Figure S2). The plot indicated a distinct group of dyads with kinship ≥ 0.07 that were most likely first cousins rather than being unrelated. While other cousin pairs were likely present in the dataset, we could not differentiate these from unrelated pairs.

#### 2.2.6 Analyses of egg count data

Egg count data for all four sampling sites was acquired in additional sampling in November 2019 and February 2020, using 120 and 60 oviposition traps, respectively (as described in 2.1). The data was used to investigate whether egg counts differed between sites and if they were consistent over both time points. The same researcher counted all eggs using a light microscope to ensure consistency. Additionally, we compared predicted egg counts throughout the sampling area with spatially-varying estimates of Wright’s neighbourhood size (NS: Wright 1946), calculated in in the *R* package *sGD* (Shrik & Cushman 2011).

Ordinary Kriging was performed in *R* using the *geoR* package to interpolate data on a map and visualise the pattern of egg numbers across different sampling sites. We created interpolative maps predicting egg numbers throughout the study area for both timepoints separately to test for a consistent pattern. Egg numbers were cube transformed to achieve normal distribution before fitting semivariograms with different covariance models. The model returning the lowest SSQ value was chosen as the best-fitting model for the data. Results from the Kriging model were cross-validated using the *xvalid* function, which compares observed values with those predicted by kriging. We visualised the results through the *image* function. Eggs counts were back-transformed before being plotted onto the map.

We used *sGD* to estimate spatially explicit indices of NS, treating our dataset as a continuous population isolated by distance (Shirk & Cushman 2014). NS represents the effective number of *Ae. notoscriptus* that contribute to the local breeding ‘neighbourhood’ under isolation by distance. Wright (1946) defined a genetic neighbourhood as NS=4πσ^2^D, which connects parent-offspring dispersal (σ) to the effective density (D) of breeding adults in the local area. The relationship between NS and D is thus highly correlated, with population density increasing linearly with NS (Sumner *et al*. 2001). We therefore used the covariation in NS estimates to assess whether egg counts could be used as a predictor for the population density of mosquitoes in the area.

We determined the genetic neighbourhood diameter using the distance class that showed the most significant positive genetic correlation calculated in the *mgram* function described above (i.e., 1300 m; radius = 650 m). We set the minimum population size to 20 individuals to minimise sampling error. We used the defined local neighbourhood around each sampling site to interpolate NS throughout the study area. Ordinary Kriging was performed in R using the *geoR* package to interpolate data on a map and visualise the pattern of NS across the sampling area. We fitted several semivariograms with different covariance models, and the model returning the lowest SSQ value was chosen as providing the best fit. Results returned by the Kriging model were cross-validated using the xvalid function. We visualised results through the *image* function. In addition, we estimated NS for the entire dataset, using the inverse of the regression slope of pairwise individual genetic distance (Rousset’s *a*) against the natural logarithm of geographical distance (Rousset 2000). We omitted pairs <100m apart as the linear relationship of genetic and geographical distance may break down at distances within the dispersal estimate σ.

## 3. Results

### 3.1 Population structure and dispersal

#### 3.1.2 Genetic diversity and local population structure

Pairwise F_st_ estimates were lowest between the ‘north-east’ and ‘south-east’ sites and highest between the ‘north-west’ and ‘south-east’ sites (Table 1; Figure 1). All F_st_ estimates were very low.

**Table 1:** F_st_ estimates between sampling sites. All standard errors = 0.00000. Site locations are visualised in Figure 1.

|  | North-west | Central | South-east | North-east |
| --- | --- | --- | --- | --- |
| North-west |  | 0.00163 | 0.00165 | 0.0013 |
| Central |  |  | 0.0008 | 0.0008 |
| South-east |  |  |  | 0.0005 |

The *fineRADstructure* plot (Figure 2) showed no clear clustering of coancestry between mosquito pairs collected from the same sampling sites, indicating gene flow between all sites.

**Figure 2.**
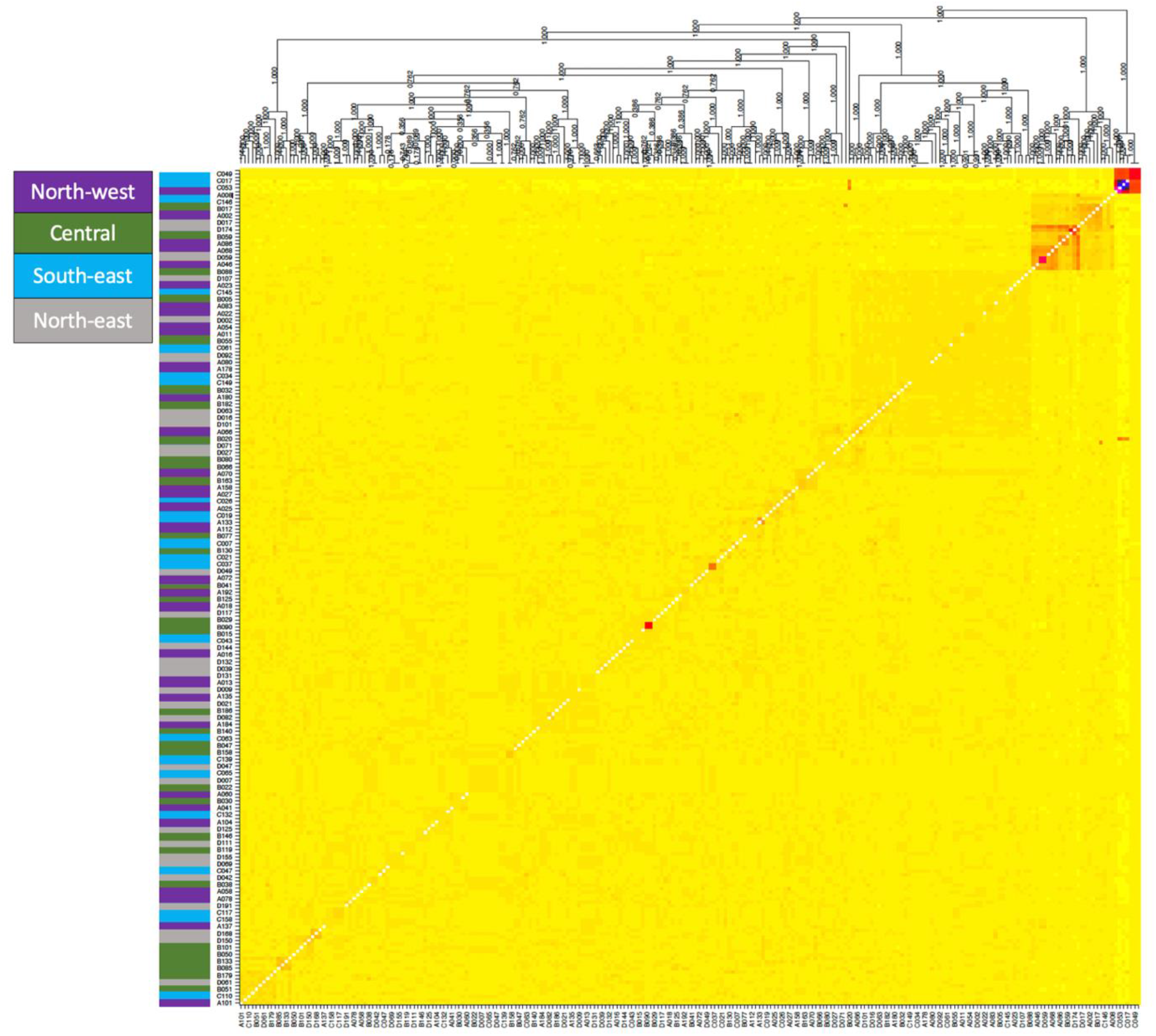
fineRADstructure coancestry map and tree. The left-hand side panel indicates genotype sampling site (‘north-west’ in magenta; ‘central’ in green; ‘south-east’ in light blue; ‘north-east’ in grey). The central panel shows co-ancestry between genotypes, with light yellow indicating low co-ancestry, and darker yellows, oranges and reds indicating progressively higher co-ancestry. The tree at the top shows inferred relationships between samples.

We found isolation by distance (IBD) in the positive relationship between genetic distance and the natural logarithm of geographical distance when calculated on the entire *Ae. notoscriptus* dataset (Bonferroni-corrected P-value = 0.02, r = 0.05, mean distance between trap pairs = 2045 m). However, when related individuals were removed from the dataset the relationship weakened (P-value = 0.07, r = 0.09), indicating that IBD at the scale of the whole sampling area is driven by closely related individuals. No IBD was detected when tested within each sampling site, where the mean distances between trap pairs was 522 m, suggesting no clear genetic structure for *Ae. notoscriptus* at this scale.

We found positive and significant spatial autocorrelation among *Ae. notoscriptus* samples around the range of 1300 m (r = 0.02, p = 0.005) as well as at 3700 m (r = 0.03, p = 0.02) (Figure 3A). At distances ranging from 4700 m onwards, *Ae. notoscriptus* showed significantly negative autocorrelation (r = −0.05, p<0.001) where individual mosquitoes were effectively no more related than they would be at random. By comparison, the population of *Ae. aegypti* from Cairns showed a signature of spatial structure that was positive and significant in the first distance class of 100 m (WT = wild type: r = 0.08, p = 0.02; wMel = *Wolbachia* infected: r = 0.05, p = 0.001), then decreased sharply, dropping below zero beyond 500 m (Figure 3B, C). In *Ae. aegypti* significant negatively autocorrelated values were observed from around 1100 m (WT: r = −0.104, p = 0.005; wMel: r = −0.05, p = 0.01). The range of trap distances for WT *Ae. aegypti* was 35 m to 3280 m and 36 m to 3057 m for the *w*Mel infected population (Figure 3B, C).

**Figure 3.**
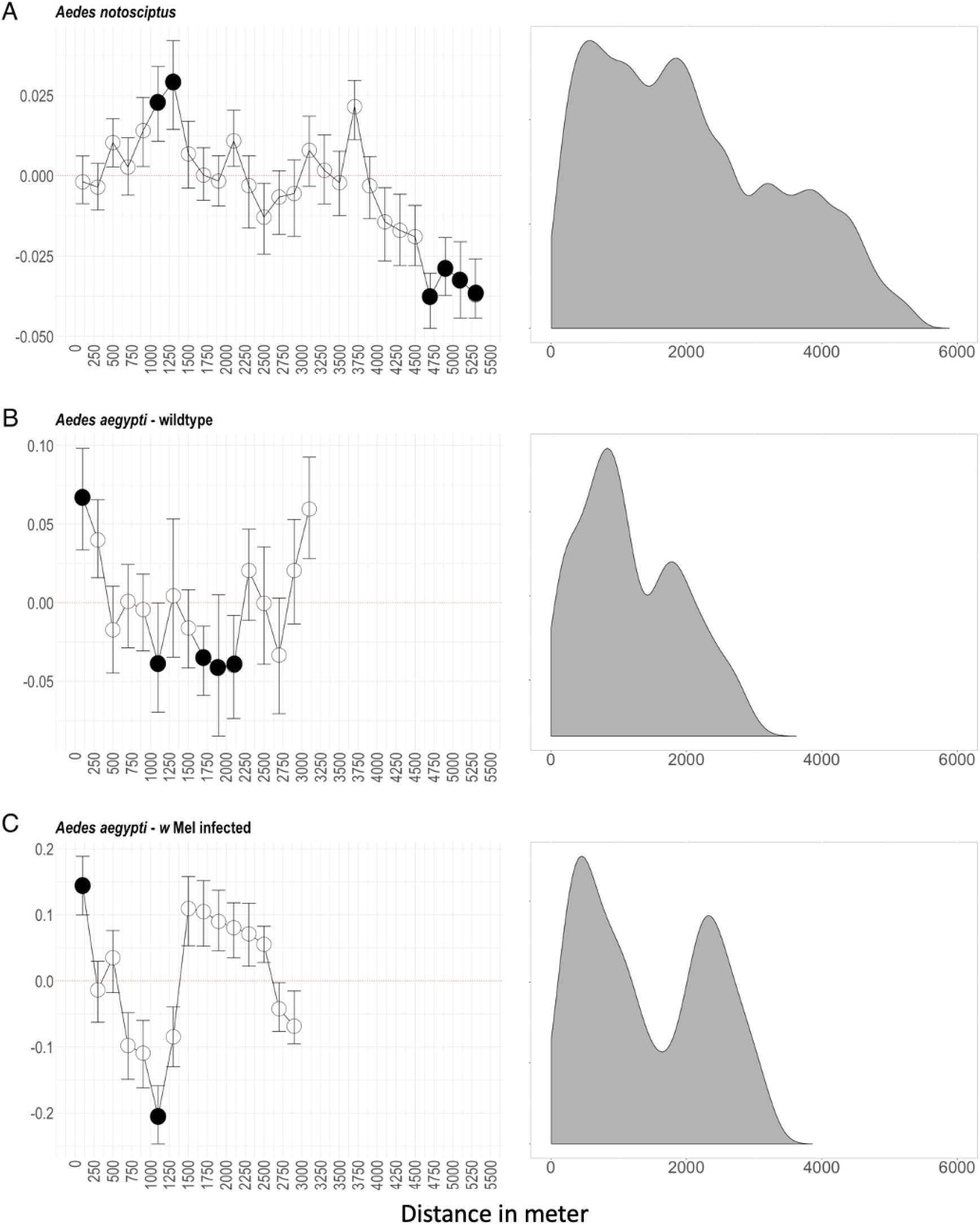
Spatial autocorrelation (left) and density of trap distances (right) of Aedes notoscriptus. **(A)**, WT (wild type = uninfected) Aedes aegypti (**B**) and wMel infected Aedes aegypti (**C**). Mantel correlograms on the left depict the association between genetic distance and geographical distance among pairs of the same distance class of 100m. Error bars show 95% confidence intervals and were calculated using 999 bootstrap replicates. Significant associations (α = 0.05) are shown as filled circles. Density graphs on the right describe the pairwise distances between traps of the three datasets. Ae. aegypti data is from Schmidt et al. (2018).

#### 3.1.2 Local dispersal estimates of individuals

We identified three putative full-sibling and 8 half-sibling pairs using *PC-relate*. We also designated 8 pairs with k > 0.07 as putative first cousins. Jasper *et al*. (2019) estimated that if the local population is constant, twice as many cousins than sibling pairs can be assumed as an average of two offspring from any one individual will themselves have offspring. However, here we present cousin pairs that we could confidently separate from unrelated individuals, meaning that cousin pairs are likely underrepresented.

Full-siblings were separated by a mean distance of 466 m (median = 179m) and exhibited a maximum observed distance of 1267 m. We found a mean separation distance for half-siblings of 1296 m (median = 340 m, max = 5173 m), and 2778 m (median = 2825 m, max = 4664 m) for first cousins (Figure 4).

**Figure 4.**
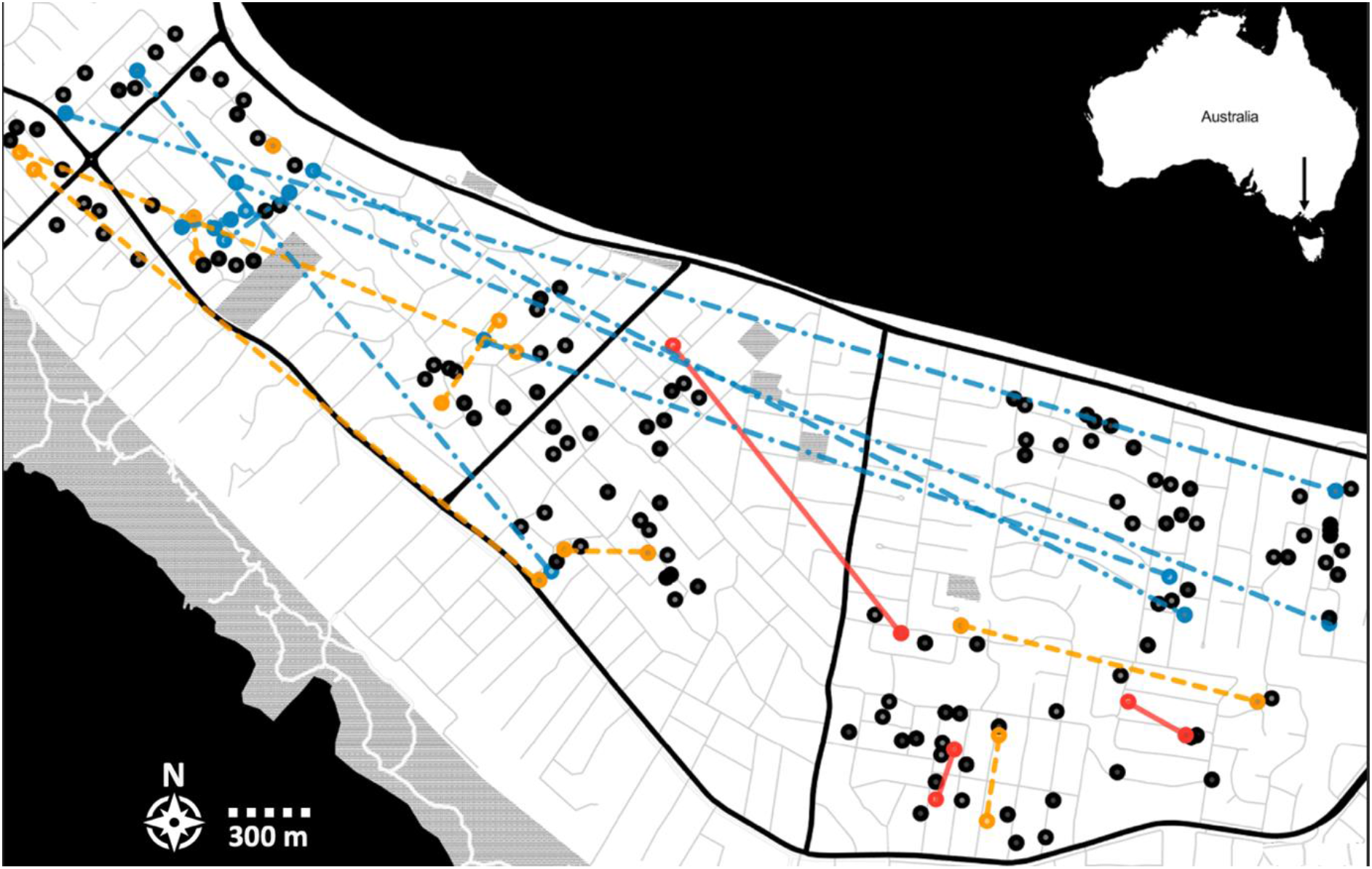
Trap placement and kinship network at the Mornington Peninsula. Circles represent trap locations. Coloured lines and circles indicate pairs of full-siblings (red, solid), half-siblings (orange, single-dashed) and first cousins (blue, double-dashed).

We detected two full-sibling pairs within the same site (‘south-east’), while the third full-sibling pair was found between the ‘ central’ and ‘ south-east’ sites (Figure 4). Most half-sibling pairs were detected within the same site (‘north-west’, ‘central’, and ‘south-east’), with one pair in the same trap one week apart and two pairs distributed between the ‘north-west’ and ‘central’ sites (Figure 4). We found pairs of first cousins between the ‘north-west’ site and the sites in the ‘north’ and ‘south-east’ with just three of the pairs within the same site (‘north-west’) (Figure 4).

### 3.2 Graphical analyses of egg counts and neighbourhood size

#### 3.2.1 Egg counts

While ordinary kriging was performed on cube-transformed egg counts, numbers per trap are visualized after back transformation to present observed egg counts per trap (Figure 5A, B). The plots show a pattern of increased egg counts from ‘east’ to the ‘north-west’ sites. The estimated prediction error after cross validation indicates that on average, an error of ~1.5 eggs can be expected at any given location. Patterns of spatial variation in egg counts were consistent across the two time points (Nov 2019 and Feb 2020), with higher egg counts being observed in the ‘north-west’ compared to the ‘north-east’ and ‘south-east’.

**Figure 5.**
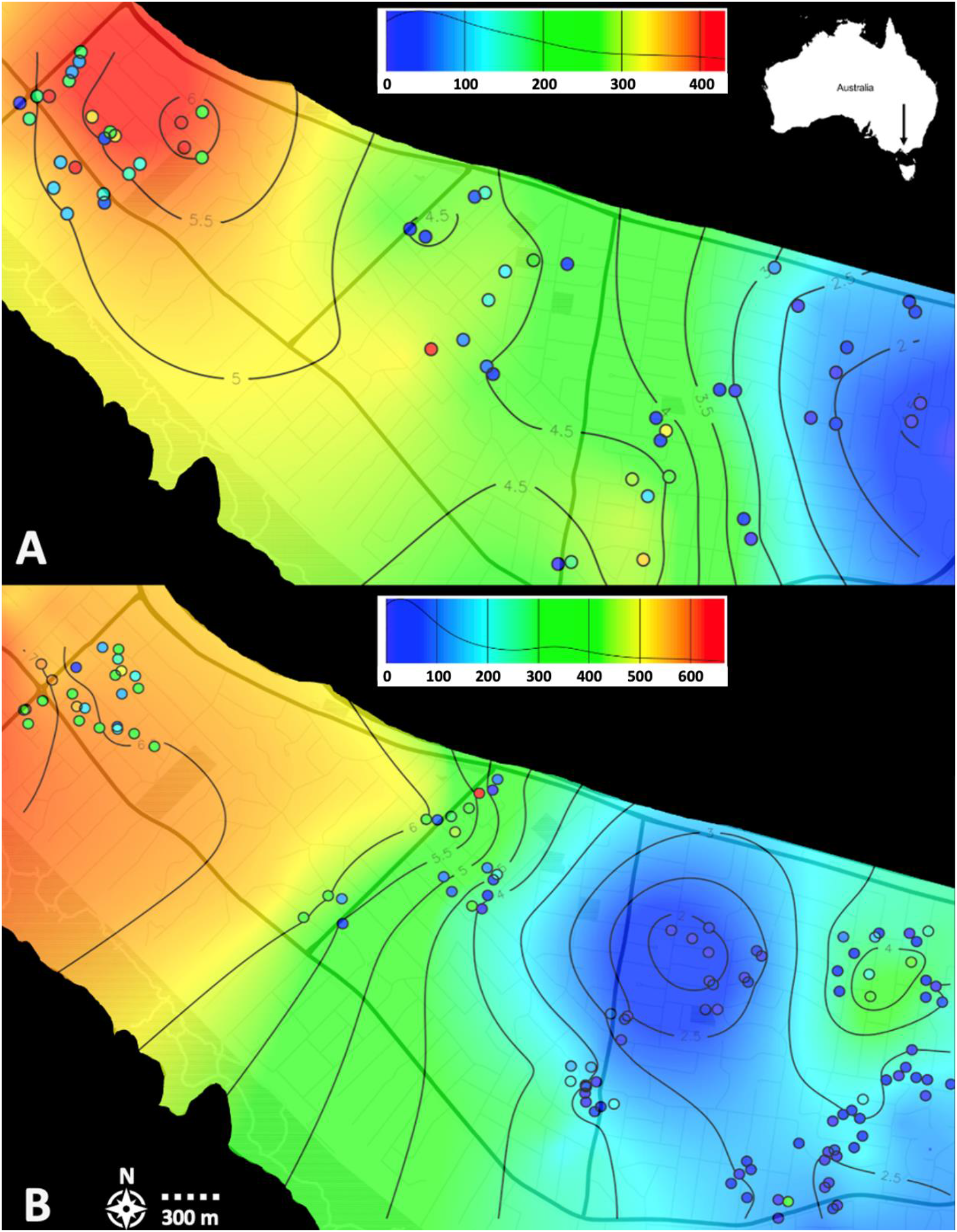
Ordinary Kriging of egg counts throughout the study area at the Mornington Peninsula. Trap locations are plotted as circles. Kriging predictions were performed on cube transformed egg counts. Colours indicate number of eggs per trap. Distribution of egg counts that kriging is based on shown in top-right panels. **(A)** Eggs collected in November 2019; **(B)** Eggs collected in February 2020.

#### 3.2.2 Neighbourhood size

The NS for each trap location estimated using *sGD* is shown as coloured circles in Figure 6, which also shows the Kriging predictions of NS throughout the area, calculated in *geoR*. The map shows lower NS (190-230) in the ‘south-east’ and ‘south-west’ sites, with an increase in NS in the ‘central’ (240-265) and ‘north-west’ (260-285) sites. The estimated prediction error for ordinary kriging after cross validation indicates that, on average, an error of ~1.04 of mean NS can be expected at any given location. Spatial variation in NS was roughly consistent with egg counts (Figures 5, 6), where higher values were observed in the ‘north-west’ and lower values in the ‘ south-east’. NS estimates produced by *sGD* were lower than using the inverse of the regression slope, with an estimated NS = 383 mosquitoes (95% C.I 287-572) across the study area.

**Figure 6.**
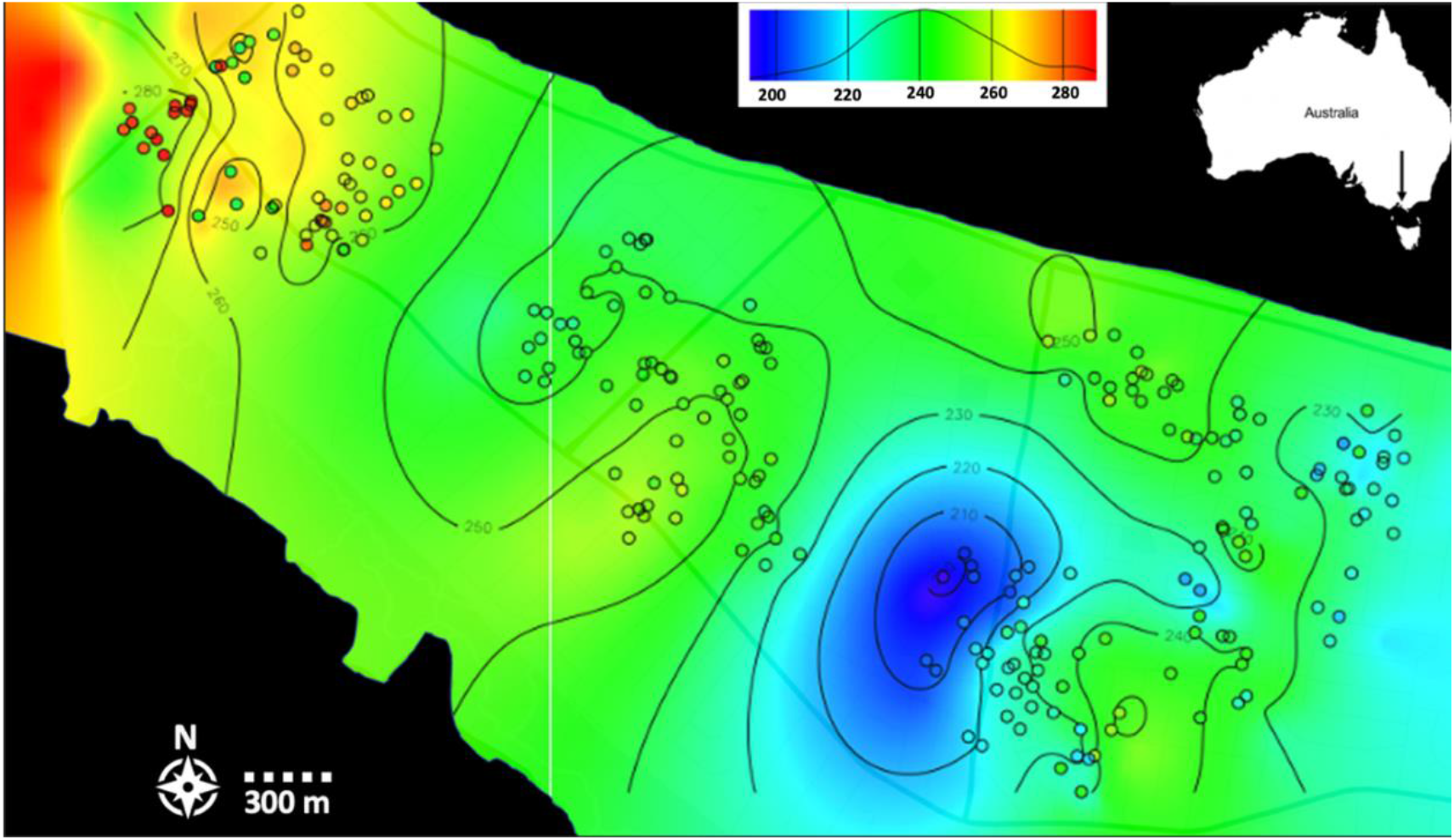
Ordinary Kriging of Neighbourhood size (NS) throughout the study area at the Mornington Peninsula. Neighbourhood size was calculated for each trap location using sGD in R. Trap locations are plotted as circles. Colours indicate NS. Distribution of NS numbers kriging is based on shown in the top-right panel. NS was calculated from genetic data from individuals collected in February 2019.

## 4. Discussion

Recent studies using spatial genomics to investigate *Aedes* mosquitoes have recorded how *Ae. aegypti* disperse within and between buildings in urban Malaysia (Jasper *et al*. 2019) and have identified dispersal barriers in *Ae. aegypti* from Saudi Arabia (Schmidt *et al*. 2022). This study used population genomic approaches to investigate the fine-scale dispersal in *Ae. notoscriptus* at the Mornington Peninsula, Australia, where the species is strongly suspected to be involved in transmitting *M. ulcerans* to humans. This represents the first application of genomic methods to *Ae. notoscriptus*, where we report both a draft genome assembly for this species and results describing how genome-wide genetic variation is spatially distributed in the Mornington Peninsula population. Our findings suggest *Ae. notoscriptus* can move greater distances than previously reported and appears to be a stronger disperser than *Ae. aegypti*. Spatial genetic variation also matched variation in egg counts, which were spatially consistent across seasons, evidence that ovitrap data may be a useful proxy for local *Ae. notoscriptus* density.

The locations of the full-sibling pairs that we detected in our kinship analysis can be interpreted as direct individual dispersal of a female mosquito that moved between the two traps during oviposition. Though we found most full-sibling pairs within the same sampling site, one full-sibling pair was detected >1km apart, showing that an individual female could disperse between two adjacent sampling sites (‘central’ and ‘south-east’; Figure 4). This distance (i.e., 1400m) is also 5-fold larger than the furthest distance reported by MRR (i.e., 238m) (Watson *et al*. 2000; Trewin *et al*. 2019). In these studies, the maximum distance of traps from the release point was 275m, so dispersal over greater distances than this would not have been detected. Conducting MRR studies over greater distances becomes far more labour-intensive, which makes close-kin methods such as in this study a valuable alternative for assessing dispersal at these scales. While we reported an average separation distance of 440m for full-siblings, it is important to note that trap placement (i.e., distances between traps) and sample selection (i.e., one individual per trap) could have impacted the detection of low-range separation distances, which would bias dispersal estimates upwards (Jasper *et al*.2022).

As in the case of full-siblings, we detected most half-sibling pairs within the same sampling site. However, some half-sibling pairs were found between adjacent sites (Figure 4), highlighting common mid-range dispersal (i.e., mean separation distance = 832m) and dispersal of individuals between sites within the same generation. We found most first cousin pairs between the sampling sites furthest apart (‘north-west’ and ‘north-east’; Figure 4), which indicates that it takes approximately two generations for *Ae. notoscriptus* to disperse through the entire study area. Overall, the results of our kinship analysis present evidence for a high dispersal activity of *Ae. notoscriptus* at the Mornington Peninsula, also supported by the low Fst values (Table 1) between sampling sites. Likewise, the lack of clustering in the coancestry analysis provided by *fineRADstructure* (Figure 2) indicates low genetic structure among sites, which reflects high gene flow between different sites, which also points towards high dispersal capacity of *Ae. notoscriptus* in the study area.

Dispersal pattern and population structure in *Aedes* mosquitoes have previously mainly focused on the dengue vector *Ae. aegypti* (e.g., Muir & Kay 1998; Harrington *et al*. 2005; Rasheed *et al*. 2013; Schmidt *et al*. 2018; Kotsakiozi *et al*. 2018; Jasper *et al*. 2019). This species’ reportedly limited dispersal has been used as a proxy for the dispersal of related container breeding mosquito species. Past MRR studies investigating dispersal in adult *Ae. notoscriptus* in urban environments of Queensland present contradictory results. Watson *et al*. (2000) describe similarly limited dispersal of *Ae. notoscriptus* compared to previous dispersal studies in *Ae. aegypti*, while Trewin *et al*. (2019) directly compared the two species in an MRR study and found that *Ae. notoscriptus* disperses further than *Ae. aegypti*. Both studies were conducted in urban areas of Queensland, with recapture trap distances ranging from 25m to 265m (Trewin *et al*. 2019) and 275m (Watson *et al*. 2000). However, both studies indicate that *Ae. notoscriptus* seems less restricted by barriers such as roads compared to *Ae. aegypti*. To further investigate differences in population structure and dispersal of the two species, we compared the results of our spatial autocorrelation analysis to two comparable datasets of *Ae. aegypti* (see 2.2.4).

Our analysis showed genetic dissimilarities between *Ae. notoscriptus* were only significant over 4km, while *Ae. aegypti* had significant dissimilarities at around 1km, which suggests greater dispersal activity in *Ae. notoscriptus* (Figure 3). Spatial autocorrelation in both the wildtype and *w*Mel infected *Ae. aegypti* revealed evidence of a strong pattern of localized genetic structure, pointing towards limited dispersal in this species. Because *Ae. notoscriptus* showed a strong positive Mantel correlation at more than tenfold larger distances than the observed correlation shown in *Ae. aegypti*, dispersal seems less restricted in *Ae. notoscriptus* and dispersal of distances >500 m may commonly occur (Figure 3). IBD detected in both *Ae. aegypti* datasets strengthens this interpretation, while we detected no IBD in *Ae. notoscriptus* at similar spatial scales (i.e., within sampling sites). We detected IBD in *Ae. notoscriptus* on the scale of the entire dataset (i.e., across all sampling sites) and calculated a neighbourhood size (NS: Wright 1946) of 287-572, which is consistent with NS calculated in *Ae. aegypti* (Jasper *et al*. 2019) and *Ae. albopictus* (Schmidt *et al*. 2021) when calculated using the inverse of the regression slope. NS calculated by *sGD* incorporated spatial variation in NS throughout the area and returned neighbourhood sizes somewhat lower.

The observed differences in dispersal behaviour of different *Aedes* species could be explained by differences in flight abilities, feeding and breeding preferences, and the likelihood of being passively moved (Verdonschot & Besse-Lototskaya 2014). While both species have adapted to breed predominantly in artificial containers, there are differences in other critical ecological factors between *Ae. aegypti* and *Ae. notoscriptus*. While *Ae. notoscriptus* does seek human hosts for blood-feeding, this species reportedly feeds on other animals such as dogs, birds, horses, possums, and fruit bats (Kay *et al*. 2008), while *Ae. aegypti* is highly anthropophilic (Harrington *et al*. 2001). This difference in host preferences may contribute to the higher dispersal activity observed in *Ae. notoscriptus*, as this species is not reliant on human blood meals to breed successfully, hence can move further away from human proximity. Additionally, MRR studies that investigated *Ae. notoscriptus* dispersal found exceptionally low male recapture rates compared to other *Aedes* species (Watson *et al*. 2000; Trewin *et al*. 2019), which could mean that male *Ae. notoscriptus* respond to different cues than other *Aedes* mosquitoes, which can influence dispersal. However, the described ecological and behavioural features of *Ae. notoscriptus* could likely differ between different clades of this species, leading to contrasting results of studies investigating traits in different populations and locations and should therefore be interpreted with caution.

Our results and analyses indicate a high dispersal ability of *Ae. notoscriptus*, which is vital to consider when planning control strategies. Trewin *et al*. (2019) argue that while *Ae. aegypti* may be controlled on the level of small blocks, the control of a highly dispersive container breeder such as *Ae. notoscriptus* will likely require much bigger areas, which is expected to be expensive and labour intensive. A pilot mosquito control study conducted in early 2021 aimed to reduce the population size of *Ae. notoscriptus* at the Mornington Peninsula non-chemically to further investigate the role of the species in the transmission of Buruli ulcer (https://www.health.vic.gov.au/infectious-diseases/beating-buruli-in-victoria; Mee *et al*. unpublished). The study employed gravid traps for four weeks to remove female mosquitoes and their offspring from the environment. The trial was evaluated by egg counts collected before and after the intervention and resulted in no measurable reduction in numbers of *Ae. notoscriptus* eggs. The success of the intervention could have been affected by several factors, including low uptake of gravid traps by residents, but was also likely affected by the size of the controlled zones. The intervention zones (~250m x 350m) were sized under the reasonable assumption that *Ae. notoscriptus* dispersal in urban areas is similar in scale to *Ae. aegypti*. However, our results highlight that generalizing ecological traits between related species can be problematic when planning species-specific interventions. The dispersal estimates we discussed above indicate that it is likely that the dispersal of *Ae. notoscriptus* exceeded the size of controlled zones. As a result, zones were likely to be invaded by mosquitoes from surrounding areas while control measures were in place, which could have compromised the success of lowering mosquito numbers. Controlled zones were also likely to be re-invaded quickly after the trial, given the generation time of *Ae. notoscriptus* of approximately one month.

Finally, the consistent egg counts observed over both time points suggests that egg counts may provide a useful indicator of localized abundance. However, local variation in egg numbers was substantial, highlighting the importance of trapping at high densities when assessing local population density with egg numbers. As applied in this study, oviposition traps represent a useful tool for this application as they are cheap, readily available, and easy to set up. We also found a trend between egg count variation and spatially explicit estimations of NS, with egg numbers and NS predictions showing lower values in the ‘north-east’ and ‘south-east’ sampling sites with increasing values through the ‘central’ to the ‘north-west’ sites at both time points (i.e., November 2019 and February 2020) (Figure 5, 6). Though NS calculations showed much less variation throughout the area than egg counts, this comparison indicates that higher egg counts correlate with larger NS and, therefore, density of mosquitoes. Our comparison of egg counts to mosquito densities matches previous studies that present a positive relationship between *Aedes* egg numbers collected by oviposition traps and numbers of adult mosquitoes collected in adult traps (Tantowijoyo *et al*. 2016; Feria-Arroyo *et al*. 2020).

### Conclusions

This study uses recently developed genomic approaches to provide evidence for high dispersal abilities of *Ae. notoscriptus* at the Mornington Peninsula, Australia. Dispersal seems to take place over greater scales in this species than in the more-thoroughly studied mosquito *Ae. aegypti*, which highlights the importance of acquiring species-specific ecological information when planning management strategies. We showed the challenge of generalizing between related species when planning interventions. Finally, we provide evidence that spatial variation in egg counts collected by oviposition traps can be linked to spatial variation in neighbourhood size and thus also density, and may be a valid means of estimating relative densities and the impact of control measures.

## Acknowledgments

We thank Nancy Endersby-Harshman and Qiong Yang for laboratory assistance and Moshe Jasper and Nancy Endersby-Harshman for helpful discussions. This research was funded by a University of Melbourne Early Career Researcher grant awarded to TLS and by the NHMRC Partnership project 1196396. VP was financially supported by the Australian Government Research Training Program Scholarship. TLS was supported by the Program Grant 1132412. AAH was supported by the NHMRC grant 1118640.

## References

1. Aguillon, S. M., J. W. Fitzpatrick, R. Bowman, S. J. Schoech, A. G. Clark, G. Coop, and N. Chen. 2017. Deconstructing isolation-by-distance: The genomic consequences of limited dispersal. PLoS genetics 13: e1006911

2. Browning, B. L., and S. R. Browning. 2016. Genotype Imputation with Millions of Reference Samples. The American Journal of Human Genetics 98: 116–126.

3. Catchen, J., P. A. Hohenlohe, S. Bassham, A. Amores, and W. A. Cresko. 2013. Stacks: an analysis tool set for population genomics. Molecular Ecology 22: 3124–3140.

4. Carvajal, T. M., K. Ogiski, S Yaegeshi, L. F. T. Hernandez, K. M. Viacrusis, H. T. Ho, D. M. Amalin, and K. Watanable. 2020. Fine-scale population genetic structure of dengue mosquito vector, *Aedes aegypti*, in metropolitan manila, Philippines. PLOS Neglected Tropical Diseases 14: e0008279.

5. Christophers, S. R. 1960. *Aedes aegypti:* the yellow fever mosquito. CUP Archive.

6. Combs, M., E. E. Puckett, J. Richardson, D. Mims, and J. Munshi-South. 2018. Spatial population genomics of the brown rat (*Rattus norvegicus*) in New York City. Molecular Ecology 27: 83–98.

7. Conomos, M. P., A. P. Reiner, B. S. Weir, and T. A. Thornton. 2016. Model-free Estimation of Recent Genetic Relatedness. The American Journal of Human Genetics 98: 127–148.

8. Danecek, P., J. K. Bonfield, J. Liddle, J. Marshall, V. Ohan, M. O. Pollard, A. Whitwham, T. Keane, S. A. McCarthy, R. M. Davies, and H. Li. 2021. Twelve years of SAMtools and BCFtools. GigaScience 10: 1–4.

9. Doak, D. F., P. C. Marino, and P. M. Kareiva. 1992. Spatial scale mediates the influence of habitat fragmentation on dispersal success: Implications for conservation. Theoretical Population Biology 41: 315–336.

10. Dobrotworsky, N. V. 1965. The mosquitoes of Victoria (Diptera, Culicidae). Melbourne University Press, London.

11. Doggett, S. L., and R. C. Russell. 1997. *Aedes notoscriptus* can transmit inland and coastal isolates of Ross River and Barmah Forest viruses from New South Wales. Arbovirus in the Asutrlian Region 7: 79–81.

12. Dray, S., and A.-B. Dufour. 2007. The ade4 Package: Implementing the Duality Diagram for Ecologists. Journal of Statistical Software 22: 1–20.

13. Edland, T. 1983. Attacks by the winter moth group (*Operophtera brumata, Agriopis aurantiaria, Erannis defoliaria*) in orchards. A system for forecasting the expected attack degree. Gartneryrket 73: 208–212.

14. Endersby, N. M., W. L. White, J. Chan, T. Hust, G. Rašić, A. Miller, and A. A. Hoffmann. 2013. Evidence of cryptic genetic lineages within *Aedes notoscriptus* (Skuse). Infection, Genetics and Evolution 18: 191–201.

15. Feria-Arroyo, T., C. Aguilar, C. Q. Vazquez, R. Santos-Luna, S. Roman-Perez, T. Oraby, G. S. Tejeda, F. C. Morales, V. S. Bueyes, and P. C. Guevara. 2020. A tale of two cities: *Aedes* Mosquito surveillance across the Texas-Mexico Border. Subtropical Agriculture and Environments 71: 12.

16. Fountain, T., A. Husby, E. Nonaka, M. F. DiLeo, J. H. Korhonen, P. Rastas, T. Schulz, M. Saastamoinen, and I. Hanski. 2018. Inferring dispersal across a fragmented landscape using reconstructed families in the *Glanville fritillar*y butterfly. Evolutionary Applications 11: 287–297.

17. Goldberg, E. E., and R. Lande. 2015. Species’ Borders and Dispersal Barriers. The American Naturalist 170: 297–304.

18. Goslee, S. C., and D. L. Urban. 2007. The ecodist Package for Dissimilarity-based Analysis of Ecological Data. Journal of Statistical Software 22: 1–19.

19. Guan, D., S. A. McCarthy, J. Wood, K. Howe, Y. Wang, and R. Durbin. 2020. Identifying and removing haplotypic duplication in primary genome assemblies. Bioinformatics, 36(9): 2896–2898.

20. Hardy, O. J., and X. Vekemans. 2002. spagedi: a versatile computer program to analyse spatial genetic structure at the individual or population levels. Molecular Ecology Notes 2: 618–620.

21. Harrington, L. C., J. D. Edman, and T. W. Scott. 2001. Why Do Female *Aedes aegypti* (Diptera: Culicidae) Feed Preferentially and Frequently on Human Blood? Journal of Medical Entomology 38: 411–422.

22. Harrington, L. C., T. W. Scott, K. Lerdthusnee, R. C. Coleman, A. Costero, G. G. Clark, J. J. Jones, S. Kitthawee, P. Kittayapong, R. Sithiprasasna, and J. D. Edman. 2005. Dispersal of the dengue vector *Aedes aegypti* within and between rural communities. The American journal of tropical medicine and hygiene, 72(2): 209–220.

23. Harris, A. F., A. R. McKemey, D. Nimmo, Z. Curtis, I. Black, S. A. Morgan, M. N. Oviedo, R. Lacroix, N. Naish, N. I. Morrison, A. Collado, J. Stevenson, S. Scaife, T. Dafa’alla, G. Fu, C. Phillips, A. Miles, N. Raduan, N. Kelly, C. Beech, C. A. Donnelly, W. D. Petrie, and L. Alphey. 2012. Successful suppression of a field mosquito population by sustained release of engineered male mosquitoes. Nature biotechnology 30: 828–830.

24. Hoffmann, A. A., B. L. Montgomery, J. Popovici, I. Iturbe-Ormaetxe, P. H. Johnson, F. Muzzi, M. Greenfield, M. Durkan, Y. S. Leong, Y. Dong, H. Cook, J. Axford, A. G. Callahan, N. Kenny, C. Omodei, E. A. McGraw, P. A. Ryan, S. A. Ritchie, M. Turelli, and S. L. O’Neill. 2011. Successful establishment of *Wolbachia* in *Aedes* populations to suppress dengue transmission. Nature 2011 476:7361 476: 454–457.

25. Honório, A. N., W. da Costa Silva, P. José Leite, J. Monteiro Gonçalves, L. Philip Lounibos, and R. Lourenço-de-Oliveira. 2003. Dispersal of *Aedes aegypti* and *Aedes albopictus* (Diptera: Culicidae) in an Urban Endemic dengue Area in the State of Rio de Janeiro, Brazil. Memórias do Instituto Oswaldo Cruz, Rio de Janeiro 98: 191–198.

26. Jasper, M., T. L. Schmidt, N. W. Ahmad, S. P. Sinkins, and A. A. Hoffmann. 2019. A genomic approach to inferring kinship reveals limited intergenerational dispersal in the yellow fever mosquito.

27. Jasper, M, T.L. Schmidt, A.A. Hoffmann 2022: Estimating dispersal using close kin dyads: The KINDISPERSE R package. Molecular Ecology Resources 22: 1200–1212.

28. Jeger, M. J. 1999. Improved understanding of dispersal in crop pest and disease management: current status and future directions. Agricultural and Forest Meteorology 97: 331–349.

29. Johnson, M. T. J., and J. Munshi-South. 2017. Evolution of life in urban environments. Science: 358.

30. Juarez, J. G., L. F. Chaves, S. M. Garcia-Luna, E. Martin, I. Badillo-Vargas, M. C. I. Medeiros, and G. L. Hamer. 2021. Variable coverage in an Autocidal Gravid Ovitrap intervention impacts efficacy of *Aedes aegypti* control. Journal of Applied Ecology 58: 2075–2086.

31. Kay, B. H., T. M. Watson, and P. A. Ryan. 2008. Definition of productive *Aedes notoscriptus* (Diptera: Culicidae) habitats in western Brisbane, and a strategy for their control. Australian Journal of Entomology 47: 142–148.

32. Kolmogorov, M., J. Yuan, Y. Lin and P. Pevzner. 2019. Assembly of Long Error-Prone Reads Using Repeat Graphs. Nature Biotechnology 37.5 540–546.

33. Kotsakiozi, P., B. R. Evans, A. Gloria-Soria, B. Kamgang, M. Mayanja, J. Lutwama, G. Le Goff, D. Ayala, C. Paupy, A. Badolo, J. Pinto, C. A. Sousa, A. D. Troco, and J. R. Powell. 2018. Population structure of a vector of human diseases: *Aedes aegypti* in its ancestral range, Africa. Ecology and Evolution 8: 7835–7848.

34. Langmead, B., and S. L. Salzberg. 2012. Fast gapped-read alignment with Bowtie 2. Nature methods 9: 357.

35. Lee, D. J., M. M. Hicks, M. Griffiths, M. L. Debenham, J. H. Bryan, R. C. Russel, and M. Geary. 1987. The Culicidae of the Australasian Region: Genus Anopheles (Anopheles, Cellia). Australian Government Publishing Service: 315pp.

36. Liew, C. C. F. C., and C. F. Curtis. 2004. Horizontal and vertical dispersal of dengue vector mosquitoes, *Aedes aegypti* and *Aedes albopictus*, in Singapore. Medical and vetenary enetomology 18.4: 351–360.

37. Malinsky, M., E. Trucchi, D. J. Lawson, and D. Falush. 2018. RADpainter and fineRADstructure: Population Inference from RADseq Data. Molecular Biology and Evolution 35: 1284–1290.

38. McCarroll, L., M. G. Paton, S. H. P. P. Karunaratne, H. T. R. Jayasuryia, K. S. P. Kalpage, and J. Hemingway. 2000. Insecticides and mosquito-borne disease. Nature 407: 6807 407: 961–962.

39. Metzger, M. E., J. W. Wekesa, S. Kluh, K. K. Fujioka, R. Saviskas, A. Arugay, N. McConnell, K. Nguyen, L. Krueger, G. M. Hacker, R. Hu, and V. L. Kramer. 2021. Detection and Establishment of *Aedes notoscriptus* (Diptera: Culicidae) Mosquitoes in Southern California, United States. Journal of Medical Entomology 59.1: 67–77.

40. Muir, L. E., and B. H. Kay. 1998. *Aedes aegypti* survival and dispersal estimated by mark-release-recapture in northern Australia. American Journal of Tropical Medicine and Hygiene 58: 277–282.

41. Palsøll, P., M. P. Zachariah, and M. Bérubé. 2010. Detecting populations in the ‘ambiguous’ zone: Kinship-based estimation of population structure at low genetic divergence. Molecular Ecology Resourses 10: 797–805.

42. Rasheed, S. B., M. Boots, A. C. Frantz, and R. K. Butlin. 2013. Population structure of the mosquito *Aedes aegypti* (*Stegomyia aegypti*) in Pakistan. Medical and Veterinary Entomology 27: 430–440.

43. Rašić, G., I Filipović, A. R. Weeks, and A. A. Hoffmann. 2014. Genome-wide SNPs lead to strong signals of geographic structure and relatedness patterns in the major arbovirus vector, *Aedes aegypti*. BMC genomics, 15(1): 1–12.

44. Reiter, P., M. A. Amador, R. A. Anderson, and G. G. Clark. 1995. Short Report: Dispersal of *Aedes aegypti* in an Urban Area after Blood Feeding as Demonstrated by Rubidium-Marked Eggs. The American Journal of Tropical Medicine and Hygiene 52: 177–179.

45. Ritchie, S. A. 2001. Effect of some animal feeds and oviposition substrates on *Aedes* oviposition in ovitraps in Cairns, Australia. Journal of the American Mosquito Contol Association 11: 2.

46. Rousset. 2000. Genetic differentiation between individuals. J Evol Biol 13: 58–62.

47. Russell, R. C., and M. J. Geary. 1992. The susceptibility of the mosquitoes *Aedes notoscriptus* and *Culex annulirostris* to infection with dog heartworm Dirofilaria immitis and their vector efficiency. Medical and Veterinary Entomology 6: 154–158.

48. Schmidt, T. L., I. Filipović, A. A. Hoffmann, and G. Rašić. 2018. Fine-scale landscape genomics helps explain the slow spatial spread of *Wolbachia* through the *Aedes aegypti* population in Cairns, Australia. Heredity 120: 386–395.

49. Schmidt, T. L., T. Swan, J. Chung, S. Karl, S. Demok, Q. Yang, M. A. Field, M. O. Muzari, G. Ehlers, M. Brugh, R. Bellwood, P. Horne, T. R. Burkot, S. Ritchie, and A. A. Hoffmann. 2021. Spatial population genomics of a recent mosquito invasion. Molecular Ecology 30:1174–1189.

50. Schmidt, T. L., Elfekih, S., Cao, L. J., Wie, S. J., Al-Fageeh, M. B., Nassar, M., Al-Malik, A., and Hoffmann, A. A. 2022. Close kin dyads indicate intergenerational dispersal and barriers. bioRxiv.

51. Shirk, A. J., and S. A. Cushman. 2011. sGD: software for estimating spatially explicit indices of genetic diversity. Molecular Ecology Resources 11: 922–934.

52. Shirk, A. J., and S. A. Cushman. 2014. Spatially-explicit estimation of Wright’s neighborhood size in continuous populations. Frontiers in Ecology and Evolution 2: 62.

53. Sumner, J., F. Rousset, A. Estoup, and C. Moritz. 2001. ‘Neighbourhood’ size, dispersal and density estimates in the prickly forest skink (*Gnypetoscincus queenslandiae*) using individual genetic and demographic methods. Molecular Ecology 10:1917–1927.

54. Sunahara, T., and M. Mogi. 2004. Searching clusters of community composition along multiple spatial scales: a case study on aquatic invertebrate communities in bamboo stumps in West Timor. Population Ecology 2004 46:2 46: 149–158.

55. Tantowijoyo, W., E. Arguni, P. Johnson, N. Budiwati, P. I. Nurhayati, I. Fitriana, S. Wardana, H. Ardiansyah, A. P. Turley, P. Ryan, S. L. O’Neill, and A. A. Hoffmann. 2016. Spatial and Temporal Variation in *Aedes aegypti* and *Aedes albopictus* (Diptera: Culicidae) Numbers in the Yogyakarta Area of Java, Indonesia, With Implications for *Wolbachia* Releases. Journal of Medical Entomology 53: 188–198.

56. Trense, D., T. L. Schmidt, Q. Yang, J. Chung, A. A. Hoffmann, and K. Fischer. 2021. Anthropogenic and natural barriers affect genetic connectivity in an Alpine butterfly. Molecular Ecology 30: 114–130.

57. Trewin, B., D. E. Pagendam, J. M. Darbro, Q. Health, and G. J. Devine. 2019. Urban Landscape Features Influence the Movement and Distribution of the Australian Container-Inhabiting Mosquito Vectors *Aedes aegypti* (Diptera: Culicidae) and *Aedes notoscriptus* (Epidemiology of Ross River virus in South East Queensland, Australia. Journal of Medical Entomology 57.2: 443–453.

58. Verdonschot, P. F. M., and A. A. Besse-Lototskaya. 2014. Flight distance of mosquitoes (Culicidae): A metadata analysis to support the management of barrier zones around rewetted and newly constructed wetlands. Limnologica 45: 69–79.

59. Wallace, J. R., K. M. Mangas, J. L. Porter, R. Marcsisin, S. J. Pidot, B. Howden, F. Omansen, W. Zeng, J. A. Axford, P. D. R. Johnston, T. P. Stinear. 2017. *Mycobacterium ulcerans* low infectious dose and mechanical transmission support insect bites and puncturing injuries in the spread of Buruli ulcer. PLoS neglected tropical diseases, 11(4): e0005553.

60. Watson, T. M., and B. H. Kay. 1999. Vector competence of *Aedes notoscriptus (*Diptera: Culicidae) for Barmah Forest virus and of this species and *Aedes aegypti* (Diptera: Culicidae) for dengue 1-4 viruses in Queensland, Australia. Journal of medical entomology 36: 508–514.

61. Watson, T. M., A. Saul, and B. H. Kay. 2000. *Aedes notoscriptus* (Diptera: Culicidae) Survival and Dispersal Estimated by Mark-Release-Recapture in Brisbane, Queensland, Australia. Journal of Medical Entomology 37: 380–384.

62. Wright, S. 1946. Isolation by Distance under Diverse Systems of Mating. Genetics 31: 39.

63. Ye, C., Ma, Z. S., Cannon, C. H., Pop, M., and W. V. Douglas. 2012. Exploiting sparseness in *de novo* genome assembly. BMC Bioinformatics 13: 1–8.

64. Zimin A. V., Marçais, G., Puiu, D., Roberts, M., Salzberg, S. L., Yorke, J. A. 2013. The MaSuRCA genome assembler. Bioinformatics 21: 2669–77.

